# Yoda molecules agonize PIEZO2

**DOI:** 10.64898/2026.05.08.723777

**Authors:** Tharaka D. Wijerathne, Aneesh Chandrasekharan, Aashish Bhatt, Yun L. Luo, Jérôme J. Lacroix

**Affiliations:** Department of Biomedical Sciences, Western University of Health Sciences, Pomona, CA, USA; Department of Biotechnology and Pharmaceutical Sciences, Western University of Health Sciences, Pomona, CA, USA

**Keywords:** Yoda1, Yoda2, PIEZO2, PIEZO1, mechanotransduction

## Abstract

Mammalian PIEZO proteins (PIEZO1 and PIEZO2) act as essential mechanosensitive channels. PIEZO1 is thought to be selectively activated by Yoda molecules (Yoda1 and Yoda2). Although a molecular mechanism for PIEZO1 activation by Yoda1 exists, the origin of Yoda1 selective agonism remains obscure. Here, using electrophysiology and calcium imaging in transfected cells, we show that Yoda1 agonizes PIEZO2 but with a distinct efficacy profile compared to PIEZO1. Yoda1 increases single channel open probability and macroscopic stretch sensitivity of PIEZO2 currents as efficaciously as reported for PIEZO1, but it elicits weak PIEZO2-dependent calcium influx in absence of external stimulus, an effect detectable only under non-physiological conditions. Yoda1 slows down inactivation of indentation-evoked PIEZO2 currents, but to a lower extent compared to PIEZO1. The Yoda1 analog Dooku1, which does not modulate PIEZO1, slows down PIEZO2 inactivation similarly to Yoda1. Two PIEZO1 agonists unrelated to Yoda1 but thought to populate the same binding site in PIEZO1 have no effect on PIEZO2 inactivation, suggesting that their PIEZO1 selectivity is better than Yoda1. PIEZO2 inactivation is also modulated by Yoda1 potent analog Yoda2. Using mutagenesis and molecular dynamics simulations, we show that Yoda2 benzoic acid group forms a transient salt bridge with a conserved arginine in the Yoda binding site (R1724/R2029 in mouse PIEZO1/PIEZO2), explaining the higher potency of this analog for both channels. Together, our study re-evaluates the PIEZO1 selectivity of Yoda molecules and highlights a complex modulatory landscape toward distinct functional properties of PIEZO channels.

## Introduction

PIEZO proteins encompass a small family of mechanically-activated and cation-selective ion channels exclusively present in eukaryotic organisms, with only two vertebrate members called PIEZO1 and PIEZO2 (1, 2). PIEZO channels share the same homotrimeric organization, that of a central ion conducting pore surrounded by three curved blade-like domains, each of these blades consisting of a juxtaposition of nine transmembrane helical bundles (called repeats A-I, or transmembrane helical units 9-1) (3-6). The inherent curvature of the PIEZO blades gives these proteins an overall bowl shape that deforms the surrounding membrane beyond the channel-membrane boundary (7-9). Mechanical membrane deformations such as lateral stretch or outward bending promote flattening of the PIEZO blades (10, 11), a conformational change thought to promote opening of the central pore (12-15).

PIEZO1 is expressed in many cell types, whereas PIEZO2 is mainly present in peripheral mechanosensory neurons as well as a discrete number of non-neuronal cells such as muscle and vascular endothelial cells (2, 16, 17). Genetic studies in which *Piezo1* and/or *Piezo2* genes have been invalidated globally or in specific tissues have revealed numerous biological roles mediated by PIEZO channels, ranging from somatovisceral mechanosensation (18-21) and proprioception (22, 23), to the growth and development of musculo-skeletal (24, 25), cardiac (26), lymphatic (27, 28) and vascular tissues (29, 30). To date, PIEZO channels have been associated with numerous ailments including blood disorders (31-34), mechanical allodynia (35-37), generalized lymphatic dysplasia (38, 39), Prune Belly syndrome (40) and abnormal musculoskeletal development (41-47).

The considerable impact of PIEZO channels on health and diseases has sparked the search for homolog-selective pharmacological modulators, both as molecular probes to dissect homolog-specific biological functions and as potential drugs to treat PIEZO-related pathologies. In this context, Yoda1, a synthetic small molecule proposed to act as a PIEZO1-specific agonist, has been discovered from a high-throughput screening campaign (48). Since its discovery, Yoda1 and its more potent analog Yoda2 (49) have been extensively used as a tool to unravel novel PIEZO1 functions in immune cells (50), bones (51-53), cancer progression (54, 55), and in the vascular system (56-59). In contrast to the common belief that Yoda1 and Yoda2 molecules act as highly PIEZO1-selective modulators, we show here that they can also agonize PIEZO2. These results show the relative promiscuity of these compounds and may prompt reinterpretation of some prior studies.

## Results

Yoda1 exerts distinct measurable effects when used at micromolar concentrations to PIEZO1-expressing cells: 1) it evokes spontaneous (i.e., without external mechanical stimulation) calcium influx; 2) it reduces the minimal force (activation threshold) required to measure ionic currents evoked by gigaseal pressurization (stretch currents) or by cellular indentations (poking currents); 3) it slows down inactivation and deactivation kinetics of macroscopic currents; and 4) it increases single channel open probability under steady-state membrane stretch conditions (48, 60, 61). The prevailing notion that Yoda1 acts as a PIEZO1 selective modulator mainly arose from Yoda1’s inability to elicit spontaneous calcium influx and to slow down inactivation kinetics of poking currents in PIEZO2-expressing cells (48). The effect of Yoda1 on PIEZO2 stretch currents has not been tested to our knowledge, perhaps because PIEZO2 was thought to be insensitive to membrane stretch until recently (15, 62, 63).

### Yoda1 robustly modulates PIEZO2 stretch currents

We first tested the ability of Yoda1 to modulate PIEZO2 stretch currents. Mechano-insensitive HEK293T^ΔPZ1^ cells (38) were transfected with a plasmid encoding mouse PIEZO2 and measured the next day to capture single channel currents in presence of 20 µM Yoda1 or vehicle control (0.05% dimethyl sulfoxide, or DMSO) at a constant patch pressure of -60 mmHg (**Figure 1a**). The pipette potential was held at +60 mV, establishing a membrane potential near -90 mV to increase the amplitude of single-channel events (60). Idealization of these currents reveal that Yoda1 increases PIEZO2’s mean open time from 16.6 ± 1.7 ms to 39.7 ± 5.4 ms (two-tailed Mann-Whitney U test p-value < 0.0001) while decreasing PIEZO2’s mean shut time from 383 ± 66 ms to 232 ± 43 ms (two-tailed Mann-Whitney U test p-value = 0.0006) (**Figure 1b**). Similar reciprocal microscopic effects of Yoda1 have been reported for PIEZO1 (60). Dividing the mean open time by the sum of mean open and shut times yielded open probability values that clearly increased in the presence of Yoda1, from 0.06 ± 0.01 to 0.17 ± 0.03 (Mann-Whitney U test p-value = 0.0021) (**Figure 1c**). A similar Yoda1-induced change in open probability was previously observed for PIEZO1 stretch currents (60). Dwell time analysis of single channel currents reveals that PIEZO2 populates at least three distinct open and shut states with distinct exponential lifetimes, all of them being reciprocally modulated by Yoda1 (**Figure 1d**), as previously observed for PIEZO1 (60). Two to three days after transfection, we next measured macroscopic PIEZO2 stretch currents evoked by brief suction pulses of incremental amplitude from +5 to -165 mmHg (**Figure 1e**). Fitting the relative amplitude of peak currents (I/Imax) using a Boltzmann function indicates that the presence of Yoda1 in the patch pipette correlates with a shift of the mid-point pressure (i.e, pressure needed to reach half of maximal activation, or P_1/2_) by about +20 mmHg (-55.4 ± 5.3 vs. -36 ± 4.5 mmHg, Mann-Whitney U test p-value = 0.0117), similar to the reported +15/20 mmHg shift for PIEZO1’s P_1/2_ value (48, 61). These results show overall that Yoda1 potentiates PIEZO2 stretch currents to a similar extent as reported for PIEZO1 (48, 60).

**Figure 1:**
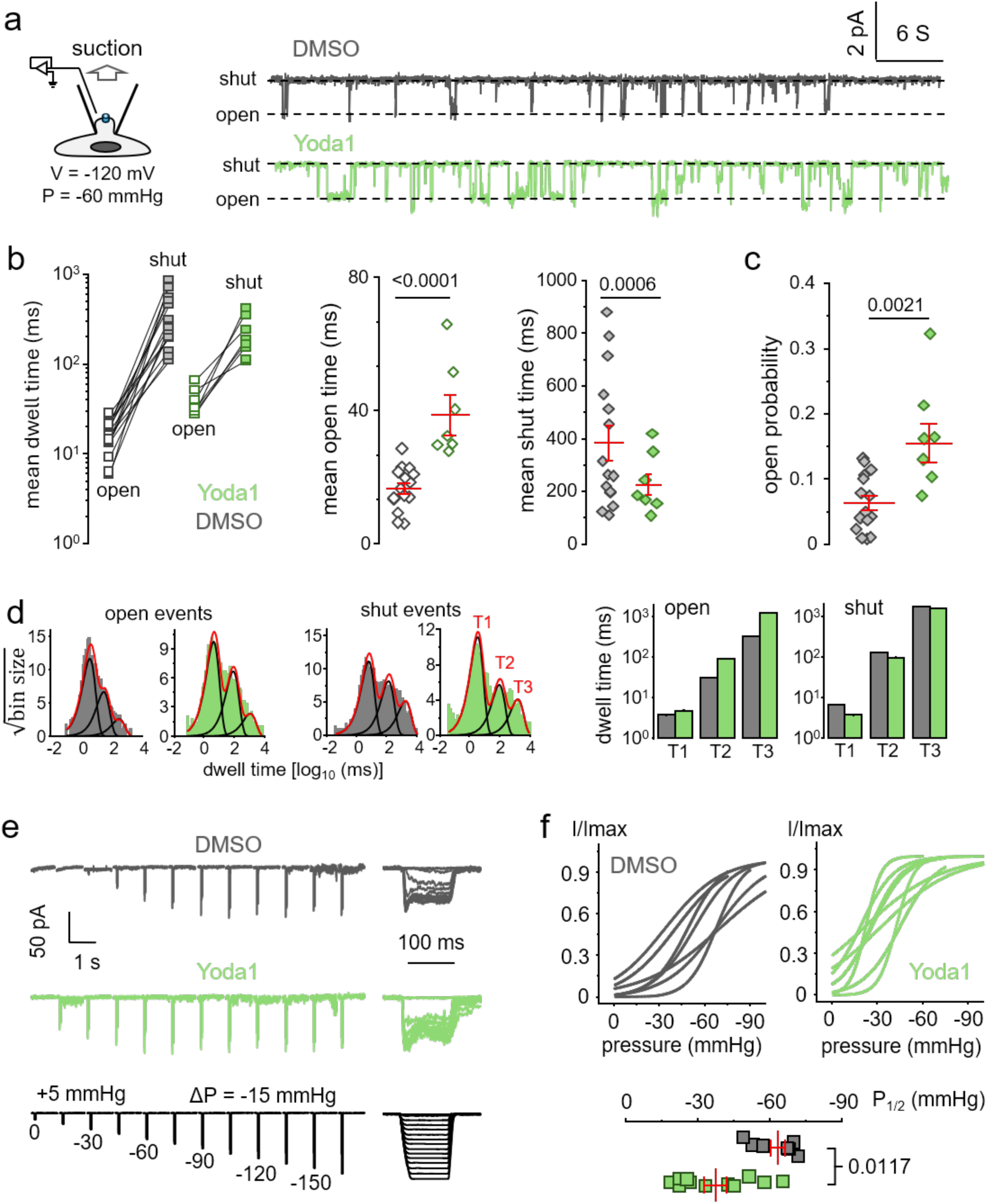
Yoda1 increases PIEZO2’s sensitivity to membrane stretch. **a**) Exemplar PIEZO2 single channel pressure-clamped current trace measured from PIEZO2-transfected HEK293T^ΔPZ1^ cells in cell-attached mode with 20 µM Yoda1 or 0.05% DMSO vehicle control in the patch pipette. **b**) Mean open and shut times from idealized single channel current traces shown in a) (DMSO: n = 15, Yoda1: n = 7 independent patches) and displayed in a paired manner for each patch (left) or unpaired to test the effect of Yoda1 (right). **c**) Open probability calculated for each patch by dividing the mean open time by the sum of mean open and shut times. **d**) *Left*: dwell time histograms obtained from single channel recordings (DMSO: 2125 shut/open events; Yoda1: 1457 shut/open events). Dwell time distributions were best fitted with three exponential components (T1-3, red traces) using Maximum likelihood estimation. *Right*: bar graphs comparing Yoda1/DMSO for each exponential component. **e**) Exemplar macroscopic PIEZO2 current traces (*left*: concatenated view; *right*: sweep stack view) evoked by brief suction pulses of varying amplitude in presence of vehicle control or 20 µM Yoda1 in the bath (*Right*: same traces shown as sweep stacks). **f**) *Top*: Graph showing relative PIEZO2 peak current (I/Imax) vs. patch pressure as idealized curves corresponding to Boltzmann fits to the data obtained from independent cells. *Bottom*: Scatter plots of the mid-point activation pressure (P_1/2_) determined from Boltzmann fits shown above (DMSO: n = 9, Yoda1: n = 12 independent cells). Error bars = s.e.m.; numbers above plots in b), c) and (f) are p-values from two-tailed Mann-Whitney U tests.

### Yoda1 can spontaneously activate PIEZO2, but only under non-physiological conditions

Given the strong modulation of PIEZO2 stretch currents by Yoda1, we sought to re-evaluate the ability of this molecule to elicit spontaneous calcium influx in PIEZO2-expressing cells, i.e, in absence of external mechanical stimulation. Two distinct scenarios could explain why this effect has not been previously observed. First, the ability of Yoda1 to modulate mechanically-evoked PIEZO2 currents (measured electrophysiologically) and to spontaneously activate PIEZO2 (measured using calcium imaging) could be mediated by distinct pharmacological mechanisms, as recently suggested for PIEZO1 (64). On the other hand, the activatory effect of Yoda1 on PIEZO2 might be too weak to be detected in absence of external mechanical stimulus, likely due to PIEZO2’s inherently higher mechanical activation threshold (15, 62).

To tease out these scenarios, we sought to test if an activatory effect can be detected by increasing the amplitude of calcium signals. To this aim, we used an isotonic “calcium boosted solution” (CBS) containing 30 mM calcium, low potassium, and no sodium, to increase the inward electrochemical driving force for calcium ions and eliminate competitive permeants (see Materials and Methods). We transfected HEK293T^ΔPZ1^ cells with a control plasmid encoding the calcium-sensitive fluorescent reporter GCaMP6m (GC6) (65), or with bicistronic plasmids encoding mouse PIEZO1 and PIEZO2 together with GC6 (61). One minute after Yoda1 perfusion, transfected cells were exposed to the calcium ionophore ionomycin to normalize maximal calcium signals (maxΔF/F_0_) as a function of the maximal GC6 response (to compensate for variability in GC6 plasmid expression).

In the presence of physiological saline (Hank’s Balanced Salt Solution, HBSS), Yoda1 failed to yield detectable PIEZO2 activation whereas it robustly activated PIEZO1 (Kruskal-Wallis with Dunn’s multiple comparison correction p-values 0.4253 vs. <0.0001, respectively) (**Figure 2a**). In contrast, in the presence of CBS, Yoda1 was able to activate both channels (Kruskal-Wallis tests with Dunn’s multiple comparisons p-values <0.0001 for both PIEZO1 and PIEZO2). We noticed that, for unclear reasons, application of DMSO alone tends to evoke small calcium transients in PIEZO2 transfected (Kruskal-Wallis with Dunn’s multiple comparison correction p-values = 0.2484 in HBSS and 0.0022 in CBS), but not PIEZO1-transfected (p-values > 0.9999) cells. This effect could be due to a weaker expression of GC6, evidenced by the overall lower absolute magnitude of ionomycin-dependent fluorescence signals in these cells (this would amplify the ionomycin-normalized amplitude of a small perfusion artefact) (**Figure 2a**). Nevertheless, because this effect is small compared to Yoda1’s, it does not invalidate the conclusion that Yoda1-induced PIEZO2 activation can be detected in absence of external mechanical stimulus, albeit under extreme, non-physiological ionic environment.

**Figure 2:**
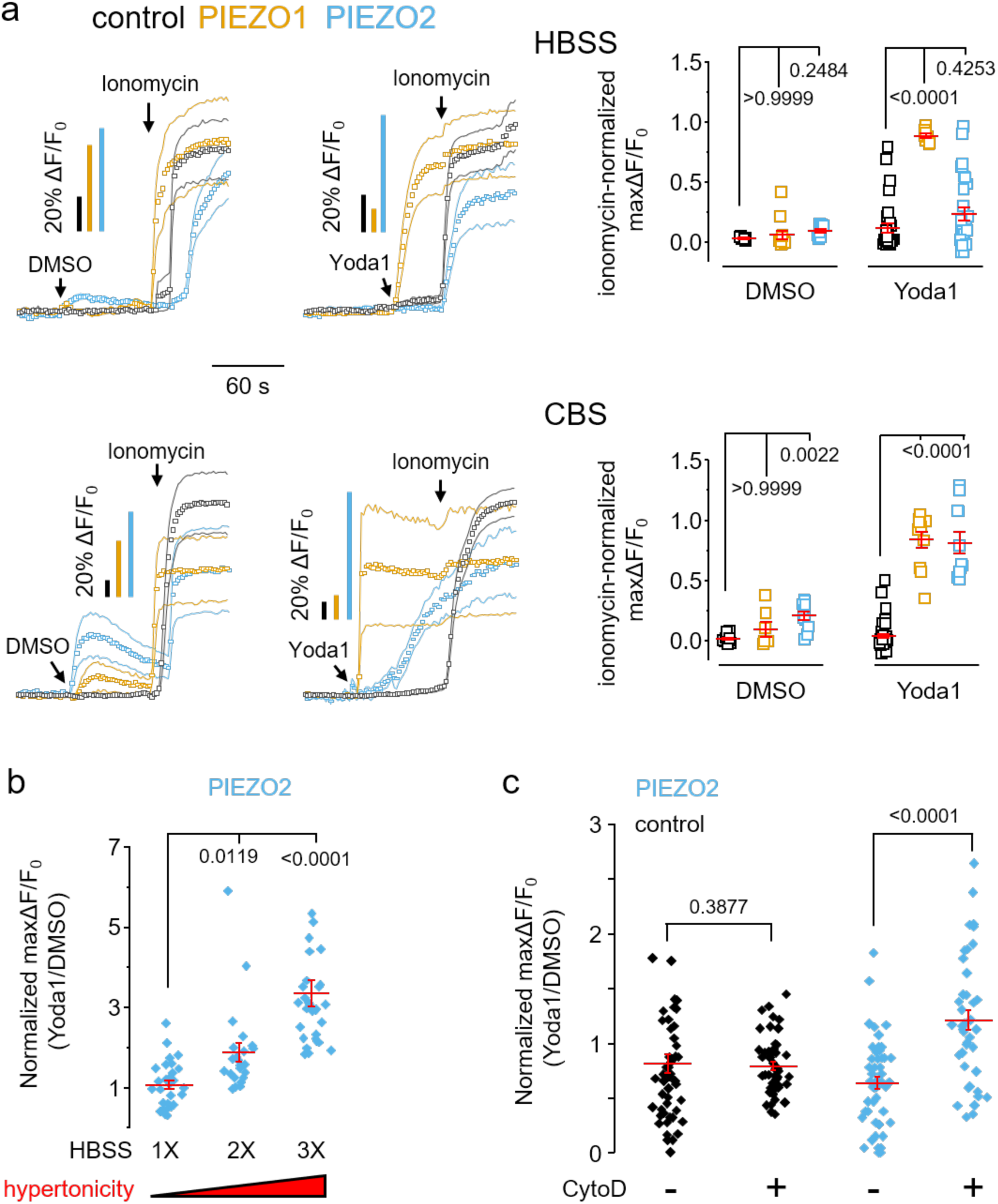
Yoda1-sensitivity of PIEZO2 probed with calcium imaging. **a**) *Left:* time-course of calcium-sensitive fluorescence obtained with confocal imaging from HEK293T^ΔPZ1^ cells transfected with GC6 only (control), with PIEZO1-IRES-GC6 (PIEZO1) or PIEZO2-IRES-GC6 (PIEZO2) plasmids in the presence of HBSS or calcium boosted (CBS) solution. Cells were first exposed to 0.075% DMSO or 30 µM Yoda1 followed by 3 µM ionomycin. Continuous lines and individual boxes respectively represent s.e.m. and mean values from n = 6 (control DMSO HBSS), 12 (PIEZO1 DMSO HBSS), 9 (PIEZO2 DMSO HBSS), 32 (control Yoda1 HBSS), 8 (PIEZO1 Yoda1 HBSS), 30 (PIEZO2 Yoda1 HBSS), 15 (control DMSO CBS), 7 (PIEZO1 DMSO CBS), 11 (PIEZO2 DMSO CBS), 48 (control Yoda1 CBS), 11 (PIEZO1 Yoda1 CBS) cells, 11 (PIEZO2 Yoda1 CBS) individual cells. *Right:* scatter plots comparison of ionomycin-normalized maxΔF/F_0_ for each condition. **b**) Epifluorescence calcium imaging from cells transfected with PIEZO2-IRES-GC6 and elicited by acute perfusion with a HBSS solution at isotonic (1X) or hypertonic (2X and 3X) concentrations containing either 30 µM Yoda1 or 0.075% DMSO vehicle (Yoda1/DMSO: 1X, n = 29/29; 2X, n = 23/18; 3X, n = 30/30 independent cells). Data were normalized by dividing maxΔF/F_0_ value for each cell treated with Yoda1 by the mean maxΔF/F_0_ value of corresponding DMSO control. **c**) Epifluorescence calcium imaging from indicated transfected cells pre-treated for 30 min with 10 µM cytochalasin D (Cyto D +: control, n = 49; PIEZO2, n = 40 independent cells) or with 0.5% DMSO vehicle control (Cyto D -: control, n = 48; PIEZO2, n = 50 independent cells) and acutely exposed to 30 µM Yoda1 or 0.075% DMSO. Data are normalized as in b). Error bars = s.e.m.; numbers above plots are p-values from (a-b) non-parametric Kruskal-Wallis test with Dunn’s multiple comparison against the indicated reference sample and (c) two-tailed Mann Whitney U tests.

The PIEZO2 structure was recently shown to adopt the characteristic flattened conformation associated with an open state (15) in response to hypertonic, but not hypotonic shocks (66). This seemed counterintuitive since PIEZO1 flattened under hypotonic, but not hypertonic conditions. This conundrum was elegantly explained by the unique property of PIEZO2 to interact with the cytoskeletal-associated protein filamin-B, altering PIEZO2’s response to distinct mechanical stimuli (66). Based on these recent findings, we next sought to test if extracellular hypertonicity and Yoda1 could act synergistically. We compared the amplitude of Yoda1-induced calcium signals (normalized to DMSO vehicle control) in cells transfected with our PIEZO2-IRES-GC6 plasmid under isotonic (1X HBSS) or hypertonic (2X and 3X HBSS) conditions. The amplitude of calcium signals were approximately 2- to 3-fold larger in hypertonic vs. isotonic conditions (Kruskal-Wallis test with Dunn’s multiple comparisons p-values = 0.0119 and <0.0001 for 2X and 3X, respectively), suggesting a larger Yoda1 activatory effect in hypertonic conditions (**Figure 2b**). Last, we destabilized actin polymerization with cytochalasin D prior addition of Yoda1. This treatment allowed us to detect a faint but significant PIEZO2-dependent activatory effect of Yoda1 in normal ionic conditions (cytochalasin D vs. vehicle: Mann-Whitney U test p-values <0.0001 for PIEZO2-expressing cells, but 0.3877 for control cells) (**Figure 2c**).

### Yoda1 modulation of PIEZO2 poking currents

Although Yoda1 was previously found unable to modulate PIEZO2 poking currents kinetics, this conclusion was based on a small sample size (48). We recorded PIEZO2 poking currents from transfected cells, either in the presence of 20 µM Yoda1 or a vehicle control (unpaired experiments). As previously observed, PIEZO2 currents display little to no inactivation when elicited by suction pulses (**Figure 1e**) (62), but robustly inactivate when elicited by indentations (**Figure 3a**) (2). When evoked with a 5 µm poke, PIEZO2 currents inactivate about 2-fold slower in the presence of Yoda1 (two tailed Mann-Whitney U test p-value < 10^-5^). For comparison, PIEZO1 poking currents inactivate 5-10-fold slower in the presence of Yoda1 under similar experimental conditions (48, 60). We did not detect a change in peak current amplitude (p-value = 0.514), nor in the channel activation threshold (non-significant differences at all tested poking distances) (**Figure 3a-b**).

**Figure 3:**
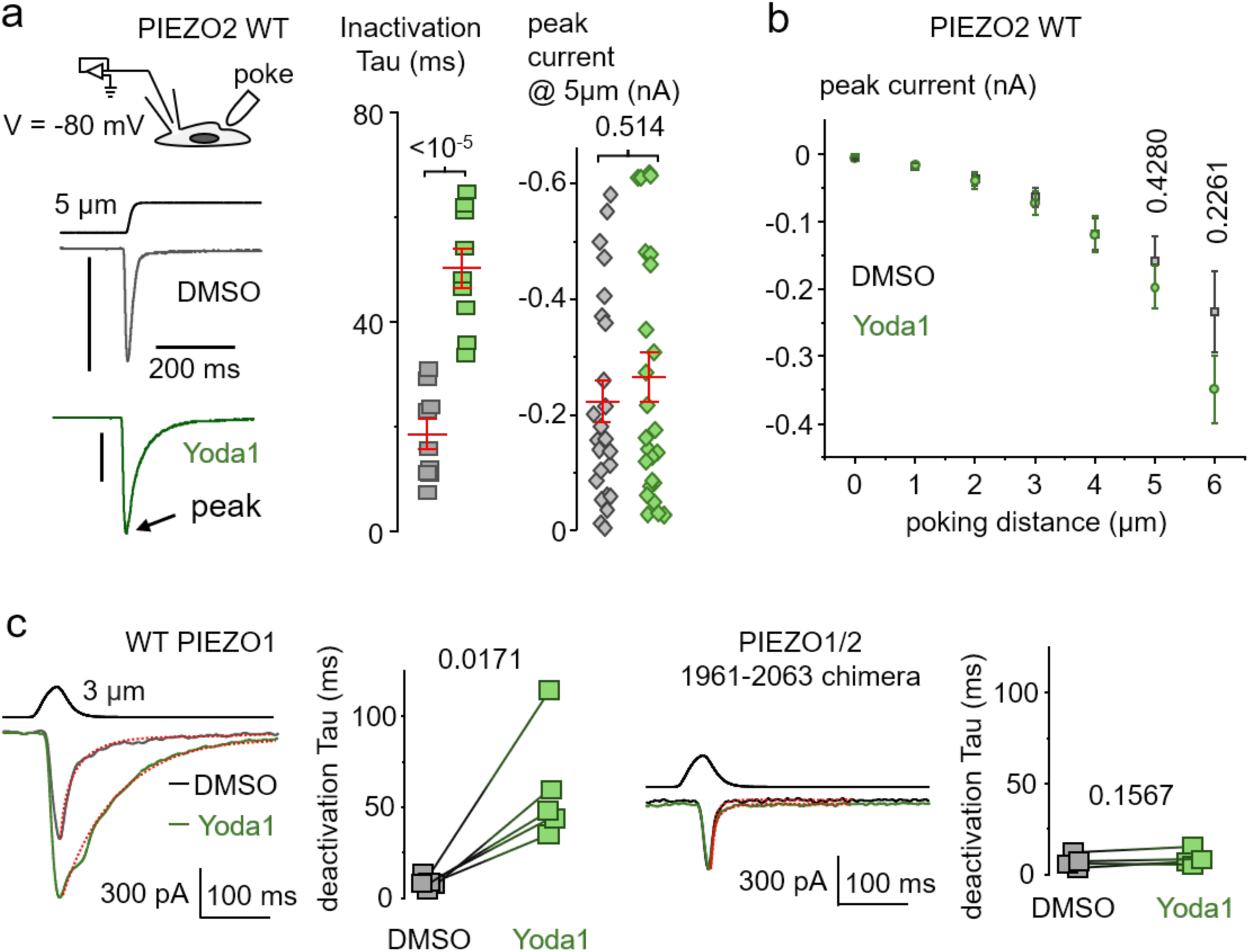
Modulation of PIEZO2 poking currents by Yoda1. **a**) *Left:* Exemplar current traces from HEK293T^ΔPZ1^ cells transfected with WT PIEZO2 in presence of 30 µM Yoda1 or 0.075% DMSO and evoked with a 5 µm poke stimulus (vertical bars represent 200 pA). *Right:* Scatter plot of inactivation Tau and peak current from traces shown on the left. **b**) Mean peak current values at various poking distances measured in both conditions described in a). **c**) Current traces evoked by a 30 ms poke stimulus in HEK293T^ΔPZ1^ cells transfected with WT PIEZO1 or the PIEZO1/2 1961-2063 chimera in the presence of 0.075% DMSO (black traces) and after addition of 30 µM Yoda1 (green traces). Next to current traces are pairwise comparison of deactivation Tau (mono-exponential fits are shown as red traces on the left) for WT (n = 5 independent cells) and chimeric (n = 4 independent cells) channels. Numbers above plots are exact p-values from unpaired t-tests with Welsch’s correction (a, b) or paired t-tests (c).

Yoda1 is thought to act as an allosteric positive modulator by binding a hydrophobic pocket sandwiched between PIEZO1 repeats A and B, promoting flattening of the mechanosensory blade domains (61, 64, 67-69). This pharmacological mechanism was initially proposed through engineering a PIEZO1/2 chimera in which a specific amino acid sequence overlapping this binding region (corresponding to mouse PIEZO1 residues 1961-2063) is swapped with its mouse PIEZO2 homologous counterpart (61). This chimera was not modulated by Yoda1 when tested using stretch current protocols, an observation which now seems odd, given the present observations that PIEZO2 stretch currents are similarly modulated by Yoda1 (**Figure 1e**). To complete our electrophysiological characterization, we sought to test an effect of Yoda1 in this chimera using a protocol giving the highest detection sensitivity. To this aim, we used a pairwise protocol with brief poke stimuli to measure deactivation kinetics, the most Yoda1-sensitive gating parameter known for PIEZO1 (60). Deactivation kinetics in this chimera is completely insensitive to 30 µM Yoda1 (p-value = 0.1567), in contrast to wild-type PIEZO1 in which it robustly slows down (paired t-test p-value = 0.0171) (**Figure 3c**). This chimera is, therefore, definitely insensitive to Yoda1.

### Modulation of PIEZO2 poking currents by other small molecules

The replacement of Yoda1’s pyrazine ring for a benzoic acid group led to the creation of Yoda2, an improved analog which agonizes PIEZO1 at nanomolar concentrations (49). We thus sought to test if Yoda2 modulates PIEZO2 currents with similar potency. Although the effect of Yoda1 on PIEZO2 is larger when probed using stretch currents, we used PIEZO2 poking currents instead because these currents are quicker to measure, facilitating the construction of dose-response curves along a range of concentrations. Using this approach, we found that Yoda2 slows down PIEZO2 inactivation kinetics at nanomolar concentrations, with a half maximal effective concentration (EC50) about one order of magnitude lower compared to Yoda1 (∼0.4 vs. ∼10 µM, **Figure 4a-b**). At saturating concentrations, however, Yoda2 exerts a smaller effect than Yoda1 (lower maximal efficacy). A similar efficacy difference between Yoda1 and Yoda2 has been reported for PIEZO1, as Yoda2 was shown to slow down PIEZO1 poking currents inactivation kinetics by 1.5-2-fold at saturating concentrations (64), while Yoda1 reportedly slows down inactivation of the same currents by 5-10 fold (**Figure 4 Supplement 1**) (60).

**Figure 4:**
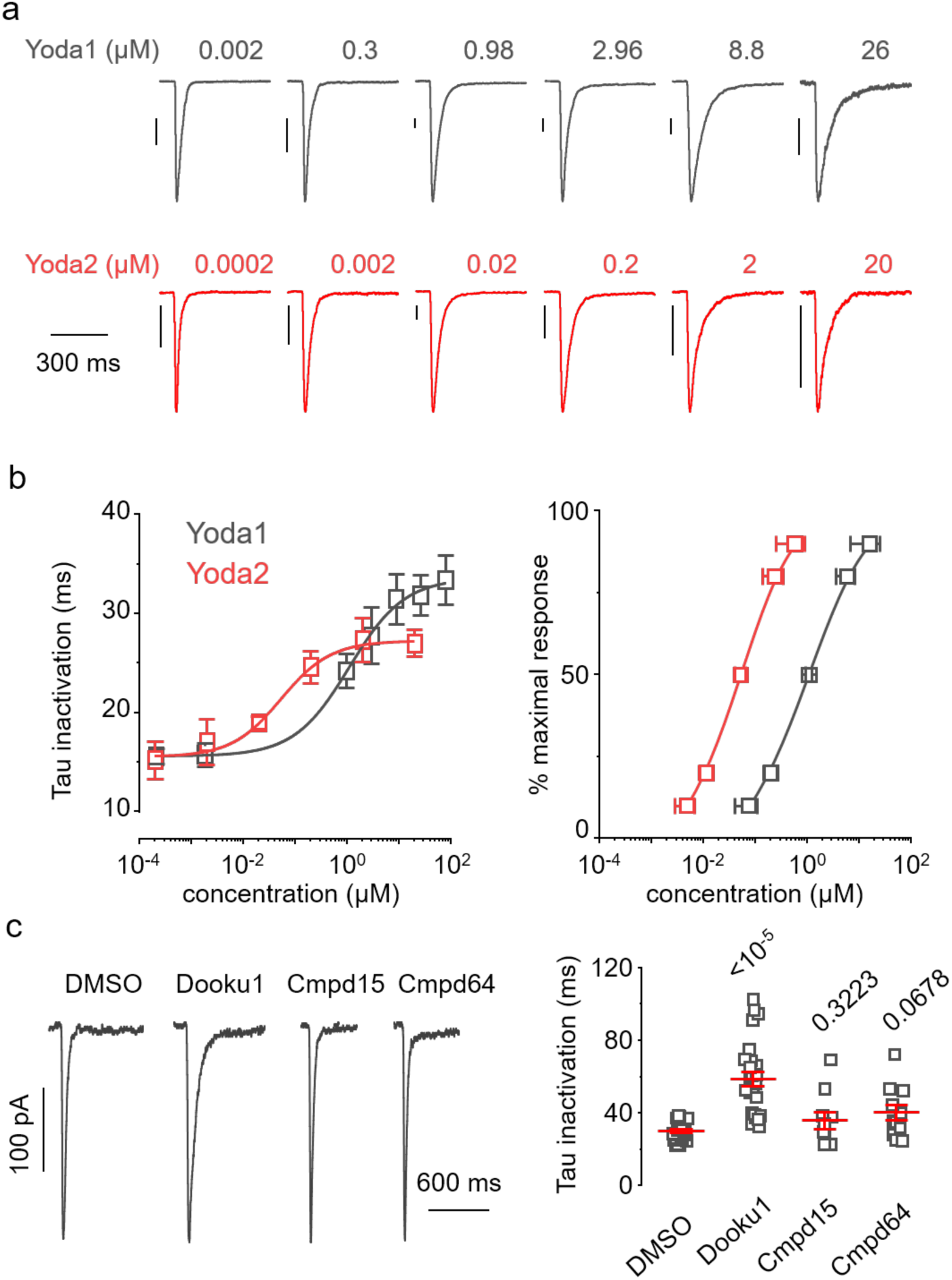
Modulation of PIEZO2 inactivation kinetics by other small molecules. **a**) Exemplar PIEZO2 poking current traces obtained at -80 mV and evoked with a 5 µm poke in the presence of Yoda1 (grey) or Yoda2 (red) in the bath solution at indicated concentrations. Vertical bars represent 200 pA for all traces. **b**) *Left*: Dose-response curves from data shown in (a) depicting mono-exponential Tau inactivation of PIEZO2 poking currents as a function of the concentration of Yoda1 (grey, n = 10, 8, 11, 11, 15, 7, 46 independent cells from low to high dose) or Yoda2 (red, n = 2, 4, 8, 10, 13, 14 46 independent cells from low to high dose) in the bath solution. Lines are fit to the data using a standard dose-response equation (see Materials and Methods). *Right*: Fitted dose-response curves normalized to the maximal response (DMSO was supplemented at all tested concentrations to match the 0.25% level present at the highest dose). **c**) *Left*: Exemplar PIEZO2 poking current traces measured in the presence of 30 µM Dooku1, Cmpd15, Cmpd64, or vehicle control (0.1 % DMSO) in the bath solution. *Right*: Scatter plots of Tau inactivation from traces shown in left (DMSO, n = 28; Dooku1, n = 27; Cmpd15, n = 8; Cmpd64, n = 8 independent cells). Numbers above plot are p-values from Kruskal-Wallis test with Dunn’s correction for multiple comparisons.

Next, we tested whether other small molecules could modulate PIEZO2 poking currents, including Dooku1 (a Yoda1 analog that antagonizes the effect of Yoda1 with no known modulatory effect on PIEZO1) and two Yoda1-unrelated PIEZO1 chemical activators (named Cmpd15 and Cmpd64) identified from a structure-guided virtual screening campaign (70, 71). Unexpectedly, Dooku1 slows down inactivation of PIEZO2 poking currents by about 2-3-fold, i.e, similar to Yoda1 (Kruskal-Wallis test with Dunn’s correction p-value <10^-5^) (**Figure 4c**). On the other hand, these currents were not significantly modulated by Cmpds 15 and 64 in our tested conditions (p-values = 0.3223 and 0.0689, respectively), suggesting that these two Yoda-unrelated activators have better PIEZO1-selectivity than Yoda molecules.

### A conserved arginine is required for Yoda2’s nanomolar potency

The Yoda binding region is well conserved between the two channel homologs, both at the level of the backbone structure and of amino acid sequence (**Figure 5a**) (3-6). It thus seems reasonable to speculate that Yoda molecules modulate PIEZO2 by binding the same region. The intracellular side of this binding region possess two conserved arginine residues, R1724 and R2098 in mouse PIEZO1, which could electrostatically interact with Yoda2’s negatively-charged benzoic acid group, potentially contributing to its higher potency relative to Yoda1 (**Figure 5b**). Although Yoda1 is electrically neutral, virtual molecular docking studies suggest that mouse PIEZO1 R2098 interacts with Yoda1’s pyrazine group (72-74).

**Figure 5:**
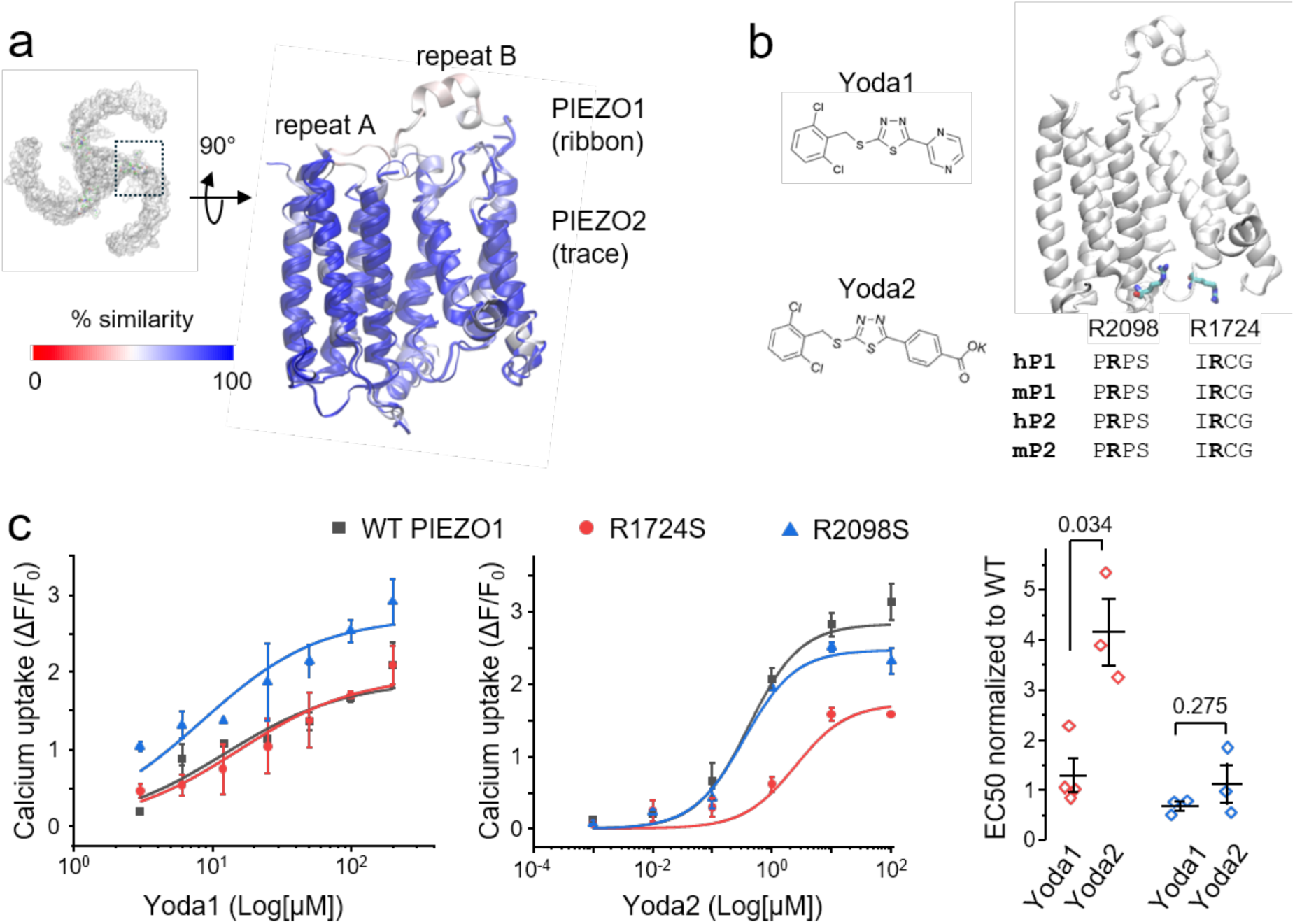
Molecular basis for Yoda2’s increased potency. **a**) Zoomed-in structural alignment of PIEZO1 (PDBID: 6b3R) and PIEZO2 (PDBID: 6kg7) showing the Yoda binding region colored as a function of sequence similarity. **b**) *Left*: chemical structures of Yoda1 and Yoda2. *Right*: localization and sequence conservation of PIEZO1 residues R1724 and R2098 (licorice). **c**) *Left*: dose-responses curves obtained for Yoda1 and Yoda2 against WT PIEZO1 (dark grey), R1724S (red) and R2098S (blue) using calcium imaging measured from n = 3 independent experiments (n = 4 for Yoda1/R1724S). Lines are fit to the data using a standard binding equation (see Materials and Methods). *Right*: scatter plots showing WT-normalized EC50 for both Yoda molecules against each mutant. The numbers next to plots indicate exact p-values from two-tailed Mann-Whitney U-tests.

We tested a pharmacological role for these basic residues in mouse PIEZO1 by substituting them for serine (to conserve their polar character while eliminating their positive charge) and evaluated the impact of these mutations on Yoda1 and Yoda2 dose-response using calcium imaging (**Figure 5c, left panels**). We fitted these dose-response curves with a standard binding equation and extracted their EC50 values. To ease comparison between mutants and WT channels, we normalized the mutants’ EC50 value to that obtained for WT PIEZO1 and plotted normalized EC50 for both Yoda1 and Yoda2 and for each mutant (**Figure 5c, right panel**). Clearly, the R2098S mutation has no effect on the potency of either Yoda1 or Yoda2, as the normalized EC50 obtained for both molecules is close to unity, ruling out a functional electrostatic interaction involving R2098. In contrast, the R1724S mutation reduces Yoda2 potency about 4-fold relative to WT, with little to no effect on Yoda1 potency (Yoda1 vs. Yoda2: two tailed Mann-Whitney U test p-value = 0.034). The large reduction of Yoda2’s potency in the R1724S mutant suggests that R1724 stabilizes Yoda2 by forming a salt bridge.

### Yoda2 forms a transient salt bridge in the Yoda binding site

The side chain of R1724 displays different orientations in different PIEZO1 experimental structures (**Figure 6a**). To quantify this orientation, we calculated the pseudo-dihedral torsion angle between the center of the guanidinium group (R1724-Cζ) and three reference atoms (**Figure 6b**). We next evaluated the temporal distribution of this torsion angle from our recent multiscale simulations of full-length PIEZO1 and PIEZO2 (15). Our analysis reveals that this side chain populates two major rotamer orientations, one with torsion angles of 50-100° and the other with torsion angles around -80°. A similar torsion angle distribution can be detected across nine PIEZO1 experimental structures (**Figure 6 Supplement 1a**). Critically, rotamers with torsion angles around -80° point the guanidinium group towards the Yoda binding site, while the other rotamers point this group away from it.

**Figure 6:**
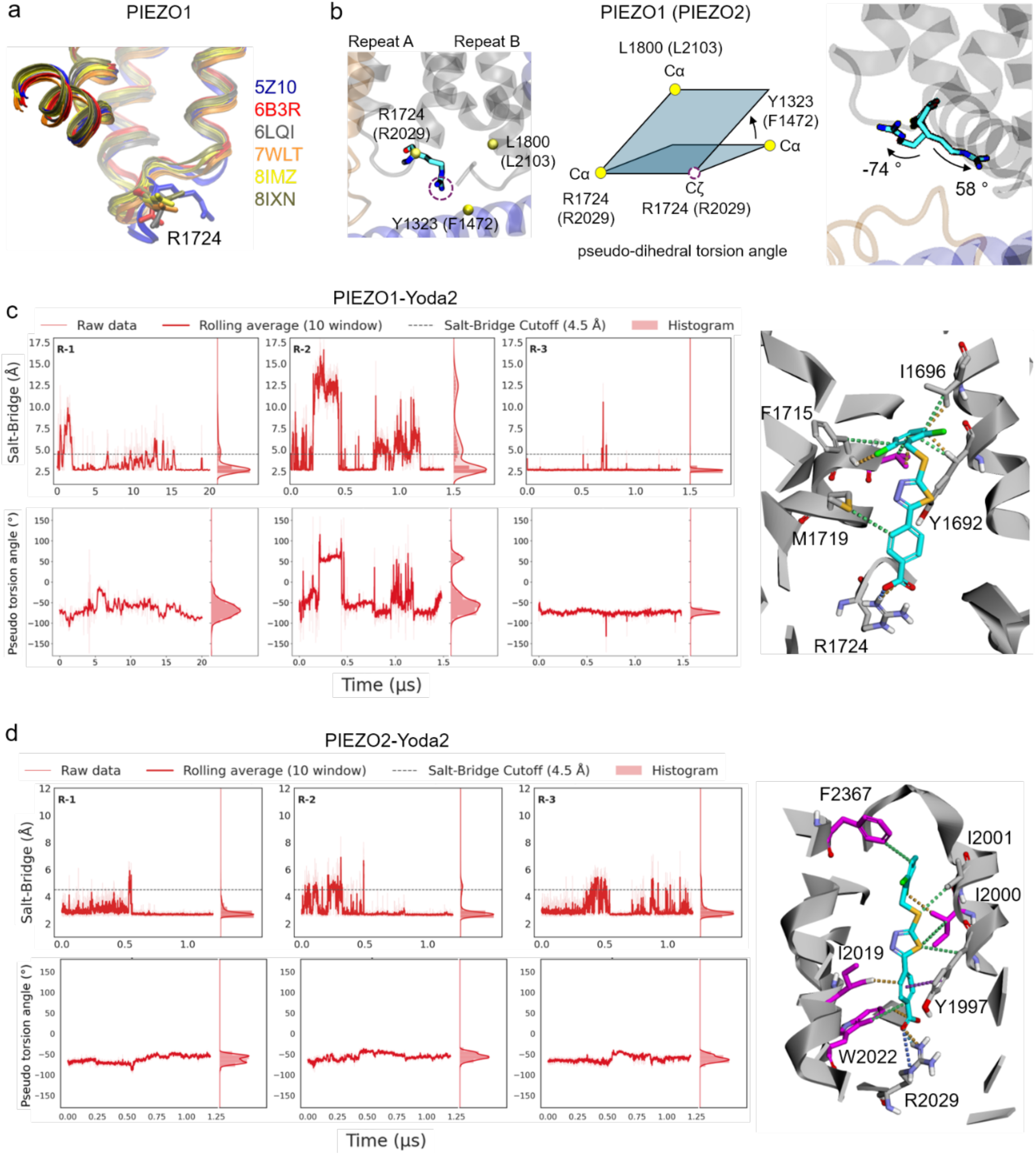
A conserved arginine electrostatically stabilizes Yoda2 in the Yoda binding site. **a**) Structural overlap of indicated PIEZO1 PDBID structures showing rotameric flexibility of R1724. **b**) *Left/middle:* Side chain orientation of R1724, and its PIEZO2 homolog R2029, calculated by measuring the angle between two planes: one formed by Cα of L1800 (L2103 in PIEZO2), Cα of R1724 (R2029) and Cζ of R1724 (R2029), the other formed by Cα of Y1323 (F1472), Cα of R1724 (R2029) and Cζ of R1724 (R2029). *Right*: example of R1724 orientations from our simulations with corresponding torsion angle. **c-d**) Three all-atom simulation replicas of a truncated PIEZO1 (c) or PIEZO2 (d) blade showing the dynamics of the Yoda2-R1724 (R2029) salt-bridge, which is predicted to form when the distance between R1724 (R2029)-Cζ and the carboxylic carbon of Yoda2 is below 4.5 Å; as well as the time-evolution of the torsion angle of R1724 (R2029). The dominant Yoda2 binding pose observed across all replicas is shown on the right for both PIEZO1 and PIEZO2 simulations.

To explore in more detail how the orientation of the guanidinium group of this conserved arginine affects Yoda2 binding, we performed three replicas of all-atom molecular dynamics (MD) simulations of a Yoda2 molecule bound to the Yoda binding site in both PIEZO1 and PIEZO2. Instead of a full-length model, we used a truncated blade (which include Repeats A-C, the anchor region and the intracellular beam) obtained from open state models for both channels. Overall, Yoda2 populates stable binding poses near the intracellular side of the Yoda binding site in both channels, similar to Yoda1 in PIEZO1 (68) (**Figure 6c-d**, **Figure 6 Supplements 1b-c and 2**). Our analysis reveals a clear correlation between the torsion angle and the formation of a salt-bridge between Yoda2 and the conserved arginine (R1724/R2029 in mouse PIEZO1/2). This bridge is stable for torsion angle values around -80° and become destabilized when R1724 or R2029 populate rotamers with positive torsion angles. In its dominant binding pose, Yoda2 also forms several non-covalent interactions with surrounding side chains in the Yoda binding site (**Figure 6c-d**).

The conserved arginine in the Yoda binding site (R1724/R2029 in mouse PIEZO1/2) also electrostatically engages with an endogenous glutamate residue (E1688/E1993 in mouse PIEZO1/2) (**Figure 6 Supplement 3).** In our previous full-length simulations, this endogenous salt bridge appears mainly in only one subunit (10 out of 13 replicas), likely because this conserved arginine is highly mobile, preventing the formation of a long-lasting bridge (**Figure 6 Supplement 4a-b**). In our all-atom simulations, the conserved arginine R1724/R2029 is closer to Yoda2’s benzoic group than E1688/E1993’s carboxy group (**Figure 6 Supplement 4c-d**), suggesting that Yoda2 is able to displace an loose endogenous salt bridge by substituting a glutamate side chain by its benzoic group.

## Discussion

Since its discovery, Yoda molecules have been used as selective PIEZO1 activators to unravel biological functions of this important mechanotransduction channel. Yet, we show here that these molecules also agonize PIEZO2, albeit with a drastically different efficacy profile. Indeed, while Yoda1 positively modulates stretch currents to a similar extent for both homologs, Yoda1 and Yoda2 only moderately slow down inactivation of PIEZO2 poking currents. Yoda1 also fails to evoke spontaneously PIEZO2-dependent calcium influx, unless the cells were osmotically stressed, their actin cytoskeleton destabilized, or their external ionic environment manipulated to favor calcium entry.

We also discovered that Dooku1, a Yoda1 analog with no effect on PIEZO1 (60, 68, 71), was able to modulate PIEZO2 inactivation of poking currents similarly to Yoda1, suggesting that this molecule could be optimized to create a PIEZO2 selective agonist. On the other hand, we show that compounds 15 and 64, two rationally identified PIEZO1 agonists (68), have no modulatory effect on PIEZO2 inactivation of poking currents. Future work will be needed to confirm whether these molecules have no modulatory effects on other functional properties of PIEZO2, in which case these molecules could represent promising PIEZO1 activators with better selectivity.

Our results further show that Yoda2 binds with a similar pose in both channels computationally captured in their flat/open conformation. In this binding pose, Yoda2’s benzoic acid group engages the guanidinium group of a conserved arginine in the Yoda binding site (R1724/R2029 in mouse PIEZO1/PIEZO2) to form an electrostatic salt bridge. We propose that the formation of this bridge, even transient, increases Yoda2’s binding affinity relative to Yoda1, enabling the detection of agonist effects at nanomolar concentrations. This work thus opens new avenues for drug design and optimization.

What molecular mechanisms could explain the different efficacy profile of Yoda molecules against PIEZO1 and PIEZO2? One possible culprit for Yoda1’s inability to evoke robust calcium influx through PIEZO2 could be its inherently high pressure-threshold for activation (62, 75). In the presence of Yoda1, PIEZO2’s activation threshold would remain higher than the resting membrane tension of the cell, limiting the frequency of spontaneous opening. Alternatively, since Yoda1 was proposed to use two distinct binding poses for spontaneous activation and current modulation of mechano-currents (64), Yoda1 may fail to occupy the proper binding pose necessary for channel activation in PIEZO2. The lower efficacy of Yoda molecules on PIEZO2 poking currents inactivation kinetics could be due to distinct interactions with cytoskeletal and adhesion proteins known to modulate inactivation kinetics of poking currents (66, 76, 77). Future studies will be needed to understand the origin of the differential pharmacological profile Yoda molecules exert on mammalian PIEZO homologs.

Our study suggests that certain roles attributed to PIEZO1 using Yoda1 *in vivo* or *in cellulo* could actually be contributed (at least partly) by PIEZO2 (especially if endogenous mechanical forces were present). For instance, reducing PIEZO2 expression in cells that co-express PIEZO1 also reduces the effect of Yoda1 (49, 78, 79). Although it is unclear whether PIEZO1 expression was reduced in these cells, this observation could be reconciled assuming a PIEZO2 contribution to the effect of Yoda1. Along the same line, since Yoda1 was shown to increase the expression of the lymphatic valve gene *ITGA9* in a PIEZO1-independent manner (27), this genetic effect could be mediated by PIEZO2, which is also present in lymphatic endothelial cells (80).

Some reported effects of Yoda1 could also be mediated by proteins unrelated to the PIEZO channel family. For example, Yoda1 was recently shown to inhibit Kv1.3 (81), a major potassium channel in immune cells. Since Yoda1 was shown to modulate certain inflammatory responses, these immunological effects could be mediated independently of PIEZO channel activation, for instance through Kv1.3 inhibition (82-84). Lastly, Yoda1 was shown to induce phosphorylation of protein kinase B (Akt) and extracellular-related kinases (ERK) 1 and 2 in vascular endothelial cells (85) in the presence of GsMTx4, a spider toxin which inhibits mechanosensitive channels including PIEZO1 and 2 (86-88).

We previously engineered a Yoda1-insensitive PIEZO1/2 chimera based on the assumption, now obsolete, that PIEZO2 is insensitive to Yoda1 (61). We show here that this chimeric swap (swapping mouse PIEZO1 residues 1961-2063 for their homologous mouse PIEZO2 counterparts) completely abolishes Yoda1’s ability to modulates channel kinetics, further strengthening the conclusion that this construct is fully insensitive to this agonist.

How could Yoda1 modulate ionic currents in both wild-type channels, but not in this chimera? In this construct, only about half of the residues forming the Yoda1 binding site (residues contained within repeat A) are swapped, implying that the chimera’s Yoda binding site is hybrid, contributed by residues from both PIEZO1 and PIEZO2. Amongst the 102 residues encompassed in this chimeric region, 21 are not conserved, meaning that this hybrid site has a different amino acid composition than in either channel. These chemical differences may allosterically modulate ligand binding affinity and/or efficacy, resulting in a weaker binding, in a binding with different, non-efficacious poses (64), or in binding poses with similar affinities between flat (open) and curved (closed) conformations, resulting in silent (non-efficacious) binding (68, 69). Further computational studies will be needed to explore these possibilities.

## Materials and Methods

### Molecular cloning

We used plasmids encoding mouse PIEZO1 (pCDNA3.1-mPIEZO1) and (PIEZO2Sport6-mPIEZO2), gifted earlier by Ardèm Patapoutian (Scripps Research). The bicistronic plasmids pCDNA3.1-mPIEZO1-IRES-GCaMP6m and pCDNA3.1-mPIEZO2-IRES-GCaMP6m were created in-house earlier (61). mPIEZO1 single point mutations were created and sequence-verified by Epoch Life Science using a pMO-mPIEZO1 plasmid.

### Cell culture, transfection and reagents

HEK293T^ΔPZ1^ cells (gifted earlier from Ardèm Patapoutian), were cultured at 37°C and 5% CO2 in Dulbecco’s Modified Eagle Medium (GIBCO) supplemented with Penicillin-Streptomycin and 5% fetal calf serum (Life technologies). HEK293T^ΔPZ1^ cells grown in 12 well plates coated with Matrigel at a 1/50 dilution (Corning), were transfected as previously described with slight modifications (89): 3 μg of mPIEZO2 plasmid or 1 μg PIEZO1 plasmid were transfected to 90% confluent HEK293T^ΔPZ1^ cells in a one well of a 12 well plate. Yoda1 (Millipore-Sigma), Yoda2 (Tocris), Dooku1 (Tocris), Cmpd15 (Chembridge chemical ID:7410103), Cmpd64 (Chembridge chemical ID:33734878), and Cytochalasin D (Millipore-Sigma) were dissolved as stock solutions in pure DMSO (Millipore-Sigma) at 40, 40, 10, 30, 30, and 2 mM, respectively. At the time of experiments, compounds were diluted to desired concentrations by dilution in the indicated solutions followed by 10-15 s vortexing. For calcium imaging experiments, compounds were extemporaneously added by manual pipetting to the cell wells.

### Electrophysiology

For single channel recordings, transfected cells were re-seeded on glass coverslips coated with Matrigel at a 1/50 dilution (Corning) one day after transfection and immediately used. Patch pipettes were pulled from G150F borosilicate capillaries (Warner Instruments) a P-97 puller (Sutter Instrument) and heat-polished using a Microforge-MF2 (Narishige), yielding a resistance of 3-5 MΩ. Single channel currents were elicited using gigaseal pressurization (high-speed pressure clamp, ALA Scientific) in HBSS (Thermofisher), acquired using an Axopatch 200B amplifier and an analog-digital converter Digidata 1550B (Molecular Devices) as individual 1 min continuous gap-free recordings using a sampling rate of 500 kHz and a filtering frequency of 5 kHz. Data were subsequently filtered at 500 Hz for display. Single channel time series were idealized using Clampfit using a dead time of 10 µs. A MATLAB (MathWorks, USA) script posted to Github (https://github.com/LacroixLaboratory) was used to ignore events with duration shorter than 0.05 ms and with change of current amplitude < 25% of the preceding event.

For macroscopic recordings, re-seeded cells were used 2-3 days after transfection. Patch pipettes were pulled to a resistance of 2-3 MΩ, and currents were acquired at 50 KHz with no post-filtering. Blunt glass probes were created by fire-polishing the tip of glass pipettes to 2-4 µm. The poking probe was angled at ∼70 ° and mounted onto a piezoelectric actuator (P-841, Physik Instrumente) controlled by Clampex *via* an amplifier (E-625, Physik Instrumente). Before each experiment, the probe was slowly moved until visual deformation of the cell. The probe was then retracted approximately by 1 µm to set the initial probe position as close as possible to the cell surface. For inactivation/deactivation measurements, a 15 second interval was maintained between stimuli to allow for complete channel recovery (60). Inactivation time constants (Tau) were calculated by fitting the decaying phase (peak to 200 ms after peak) of the macroscopic currents with a standard mono-exponential equation in Clampfit 11.2 (Molecular Devices). Dose response curves were fitted using a default dose response equation in OriginPro: Tau = 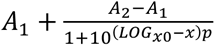; where *A*_1_ and *A*_2_ respectively represent the bottom and top asymptotic Tau values (*A*_2_ > *A*_1_); *p* represents the slope of the sigmoid curve (*p* > 0), *x* is the ligand concentration; and *LOG_x_*_0_ represents the ligand concentration at which Tau = 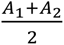

### Calcium imaging

Cells were washed and incubated for at least 30 min in HBSS prior measurements. The CBS solution contains 1 mM KCl, 30 mM CaCl2, 1 mM MgSO4, 108 mM N-Methyl-D-Glucosamine chloride, and 10 mM HEPES. The pH was adjusted to 7.4 with HCl and has an osmolarity of 288 mOsmol/L (all chemicals were purchased from Millipore-Sigma). For hypertonic treatments, 1X, 2X and 3X HBSS were prepared by dilution of a 10X commercial HBSS solution (Gibco). GCaMP6m fluorescence was recorded at 1 frame per second using a GFP/FITC filter set on an inverted Nikon Eclipse fluorescence microscope or with 488 nm laser excitation and emission detection between 500 and 550 nm on a Zeiss LSM 880 confocal microscope. Regions of interest (ROIs) were defined around individual cells, and fluorescence intensities were measured after background subtraction. Calcium responses were expressed as ΔF/F₀ = (F − F₀)/F₀, where F represents the fluorescence intensity at each time point and F₀ represents the mean baseline fluorescence intensity before compound addition. Dose-responses shown in Figure 5c were fitted using: y = 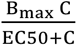; where y represents ΔF/F_0_; C represents the ligand (Yoda1/2) concentration; and Bmax represents the saturated value of ΔF/F^0^ at infinite ligand concentrations.

### Statistics

Statistical tests are indicated in figure legends and were performed using GraphPad Prism 9.

### Molecular dynamics simulations

Docking and molecular dynamics (MD) simulations used a truncated all-atom open PIEZO1 (residues 1131 to 2190) (69). For PIEZO2, we used the previously published model generated by applying membrane tension of 9.3mN/m to the structure (PDB ID: 6KG7) using the PACE force field (15). A single blade of PIEZO2 (residues 1277-2467) was retained, and the PACE model was converted to all atoms for molecular docking and MD simulations. Molecular docking was performed using AutoDock Vina 1.2.3 with the docking grid defined between Repeat B (TM29– 30) and Repeat A (TM35–36) using MGLTools-1.5.6 and a grid size of 30 × 30 × 30 Å. The PIEZO1-Yoda2 and PIEZO2-Yoda2 complex system were assembled using CHARMM36m parameters for the protein, CHARMM36 parameters for POPC lipids, ions, and water molecules, and CHARMM General Force Field (CGenFF) for Yoda2. Both systems were neutralized and maintained at an ionic concentration of 150 mM KCl and solvated using TIP3P water. The simulation system dimensions and composition are summarized in **Table 1**. All-atom MD simulations were performed using GROMACS 2024.2 smp. We followed the input files generated by CHARMM-GUI, including six equilibration steps followed by production simulations. Energy minimization was performed using the steepest descent algorithm for 5,000 steps. Long-range electrostatic interactions were calculated using the Particle Mesh Ewald (PME) method with a real-space cutoff of 1.2 nm. Van der Waals interactions were smoothly switched off between 1.0 and 1.2 nm using a force-switch function. All hydrogen-containing bonds were constrained using the LINCS algorithm, allowing a 2 fs integration timestep. The system was equilibrated at 310.15 K using the V-rescale thermostat with separate coupling groups for the protein-ligand complex, membrane, and solvent. Semi-isotropic pressure coupling was applied using the C-rescale barostat at 1 bar with a compressibility of 4.5 × 10⁻⁵ bar⁻¹ and a relaxation time constant of 5 ps. Positional restraints were applied during equilibration to the protein backbone, lipids, and dihedral angles. Following equilibration, three independent production replicas were run from distinct equilibration conformations. For PIEZO1-Yoda2, Replica 1 was simulated for 20 µs on the Anton3 supercomputer. Whereas Replica 2 and 3 were simulated for 1.4 µs each using GROMACS. For PIEZO2-Yoda2, all three independent replicas were simulated for 1.2 µs each using by GROMACS. The 20 µs replica was run on Anton3 supercomputer by converting a 100 ns GRO file to a Desmond-compatible format using Viparr on Bridges2. The Anton3 simulation was performed using a multigrator integration scheme with a 2.5 fs timestep. Temperature was maintained using a Nose–Hoover thermostat at 310.15 K and pressure was controlled using the Martyna–Tuckerman–Klein (MTK) barostat with semi-isotropic coupling.Ligand–protein interaction analyses were performed using MDAnalysis and PLIP 2.1.0, and molecular visualization was carried out using VMD 1.9.3.

**Table 1:** All atom Molecular Dynamics simulation details.

| System | Membrane system<br>(xyz dimension, Å) | Number of atoms | Duration ($\mu\text{s}$) |
| --- | --- | --- | --- |
| **PIEZO1–Yoda2** | POPC (Upper/Lower=220/201)<br>(130.12 x 13.012 x 13.46) | Total Atoms= 211358<br>Protein= 11873<br>Yoda2= 33<br>Membrane= 56414<br>Water = 142755<br>K/Cl- =130/151 | Replica 1 = 20*<br>Replica 2 = 1.4<br>Replica 3 = 1.4 |
| **PIEZO2–Yoda2** | POPC (Upper/Lower=155/132)<br>(108.87 x 107.87 x 131.17) | Total Atoms = 156427<br>Protein = 11371<br>Yoda2= 33<br>Membrane= 38458<br>Water = 106365<br>K/Cl- = 96 /104 | Replica 1 = 1.2<br>Replica 2 = 1.2<br>Replica 3 = 1.2 |

\* Replica 1 was run on the Anton3. Where the other replicas run using GROMACS software.

## Acknowledgments

We thank Ardem Patapoutian (Scripps Research) and Tibor Rohacs (New Jersey Medical School, Rutgers University) for the gift of cells and plasmids. This work was supported by NIH grants GM130834 (to JJL and YLL), HL179692 (to JJL and YLL), AT013465 (to JJL), and the Anton3 allocation award MCB190044P (to JJL and YLL), supported by NIH grant GM154042 awarded to the Pittsburgh Supercomputing Center.

## Data availability

An experimental data Excel file was deposited into the open-access repository Open Science Framework (DOI: 10.17605/OSF.IO/HQG7D).

## Author Contributions

TDW, JJL, and YLL designed research; TDW, AC, AB, and JJL performed research and analyzed data; JJL wrote the paper. AC and AB contributed equally to this work.

## Competing Interest Statement

The authors declare no competing interest.

---

**Figure 4 Supplement 1:**
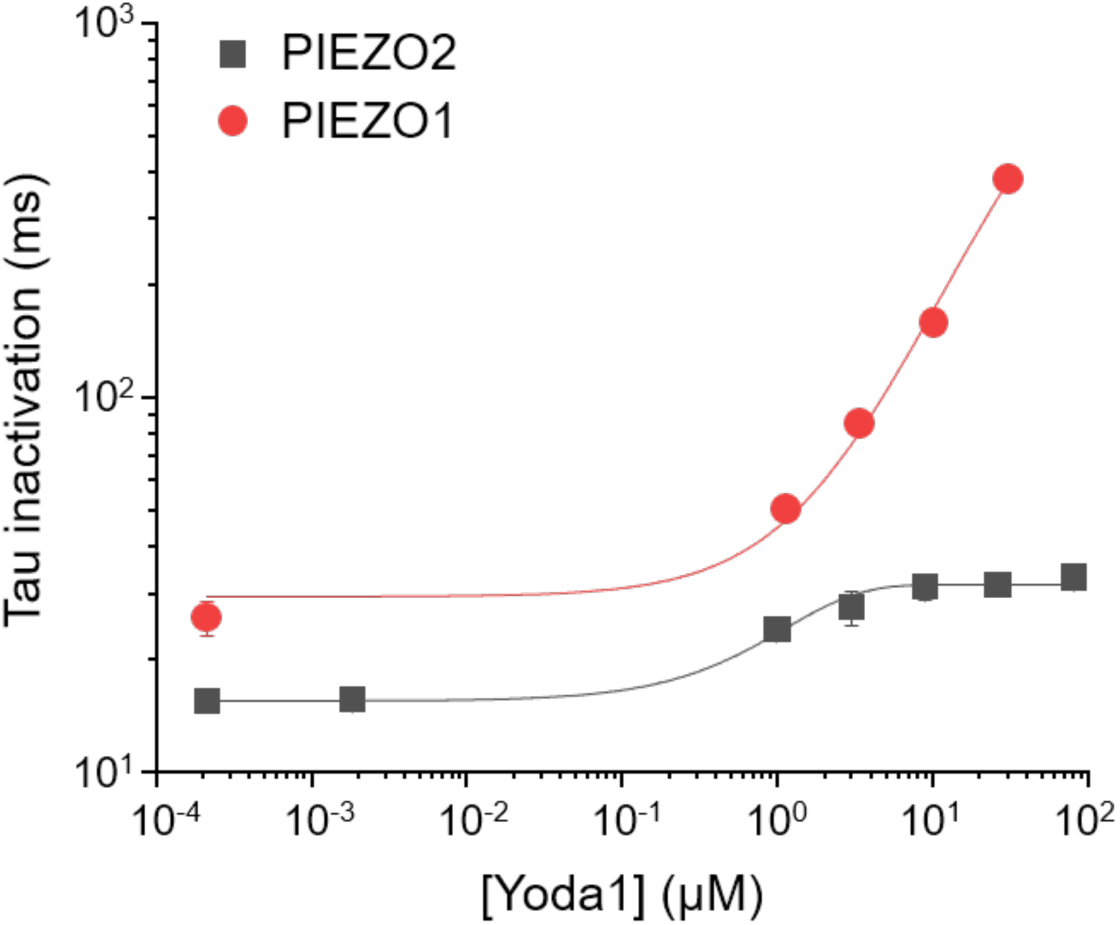
Comparison of Yoda1 efficacy between PIEZO1 and PIEZO2. The Yoda1 dose-response on PIEZO2 inactivation Tau measured in this study (grey) is compared to a previously measured Yoda1 dose-response on mouse PIEZO1 inactivation Tau (red) (60). Both axes are displayed in a logarithmic scale.

**Figure 6 - figure supplement 1:**
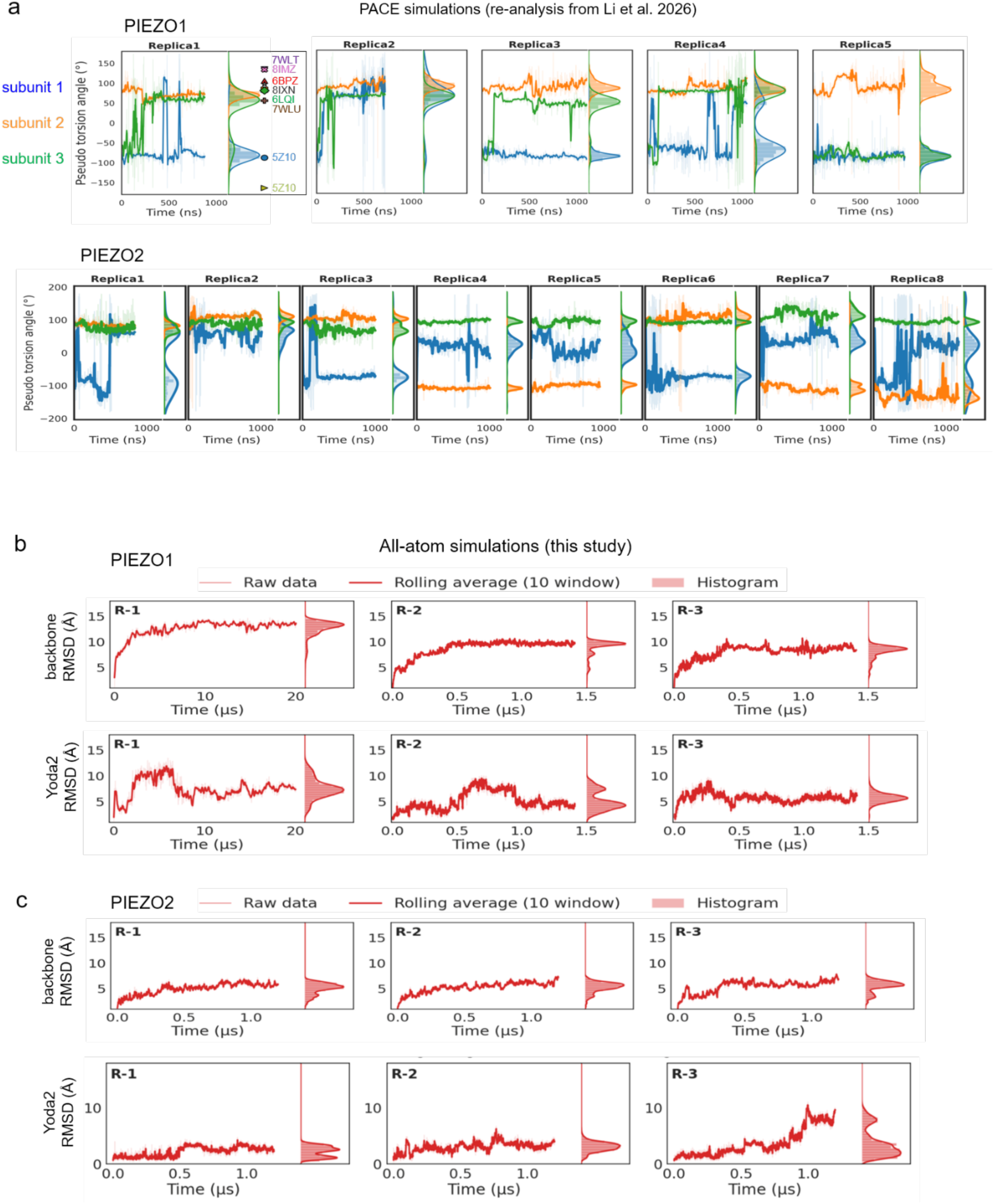
**a**) Torsion-angle dynamics of PIEZO1-R1724 (top) and PIEZO2-R2029 (bottom) from previous multiscale simulations of full-length PIEZO1 and PIEZO2 under membrane tensions of 9.0 and 9.3 mN/m, respectively, using the PACE (Protein Any-atom/Coarse-grained Parametrization with Environment) hybrid force field (15). Vertical markers adjacent to the PIEZO1-Replica1 histogram indicate the torsion angles observed in nine indicated PIEZO1 PDB structures. **b-c**) Root-mean-square-deviations (RMSD) calculated along the trajectories for the protein backbone (top) and for the Yoda2 ligand (bottom), and for both the PIEZO1-Yoda2 (b) and the PIEZO2-Yoda2 system (c).

**Figure 6 - figure supplement 2:**
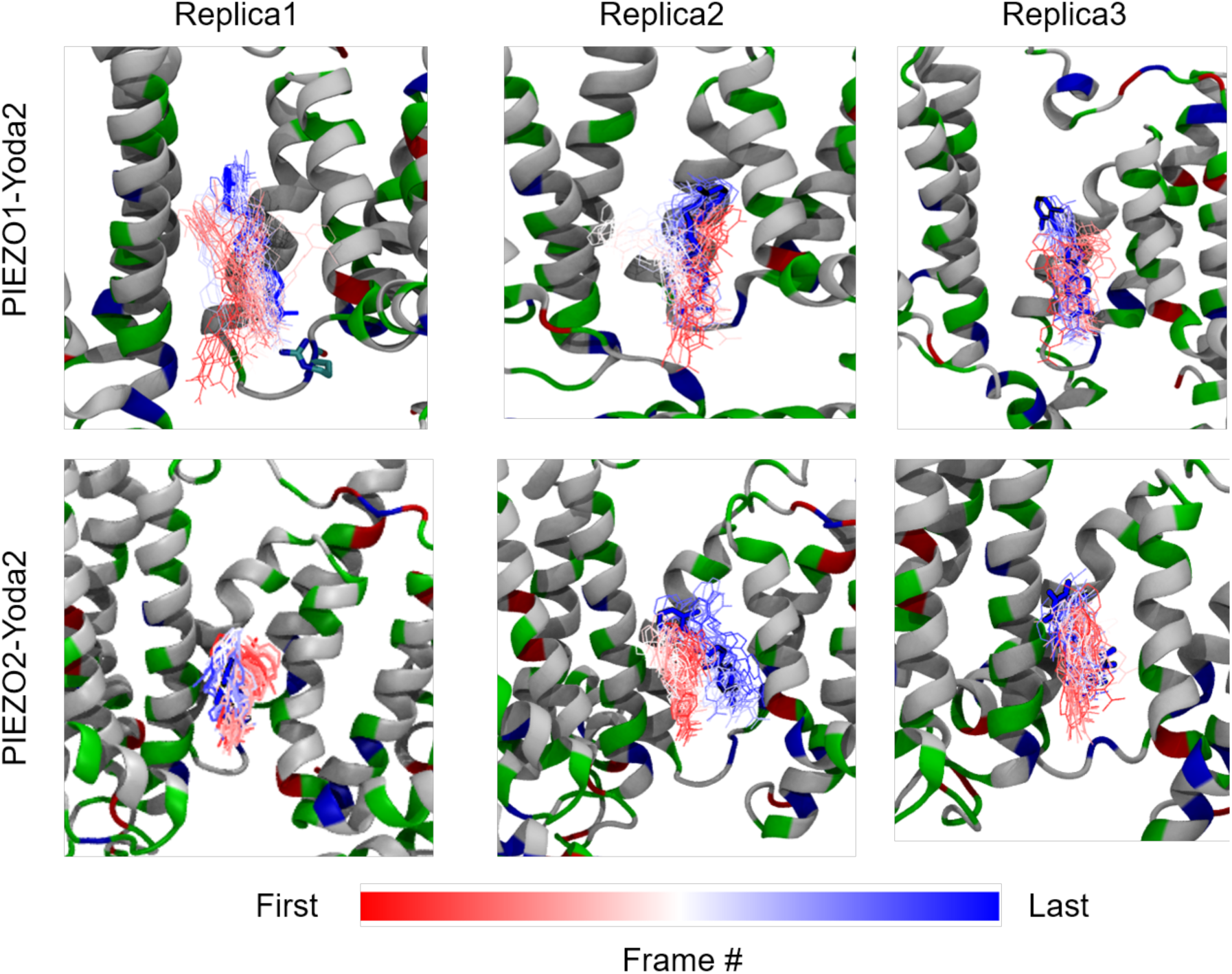
Time evolution of Yoda2 binding poses in PIEZO1 (top) and PIEZO2 (bottom) across all simulation replicas

**Figure 6 - figure supplement 3:**
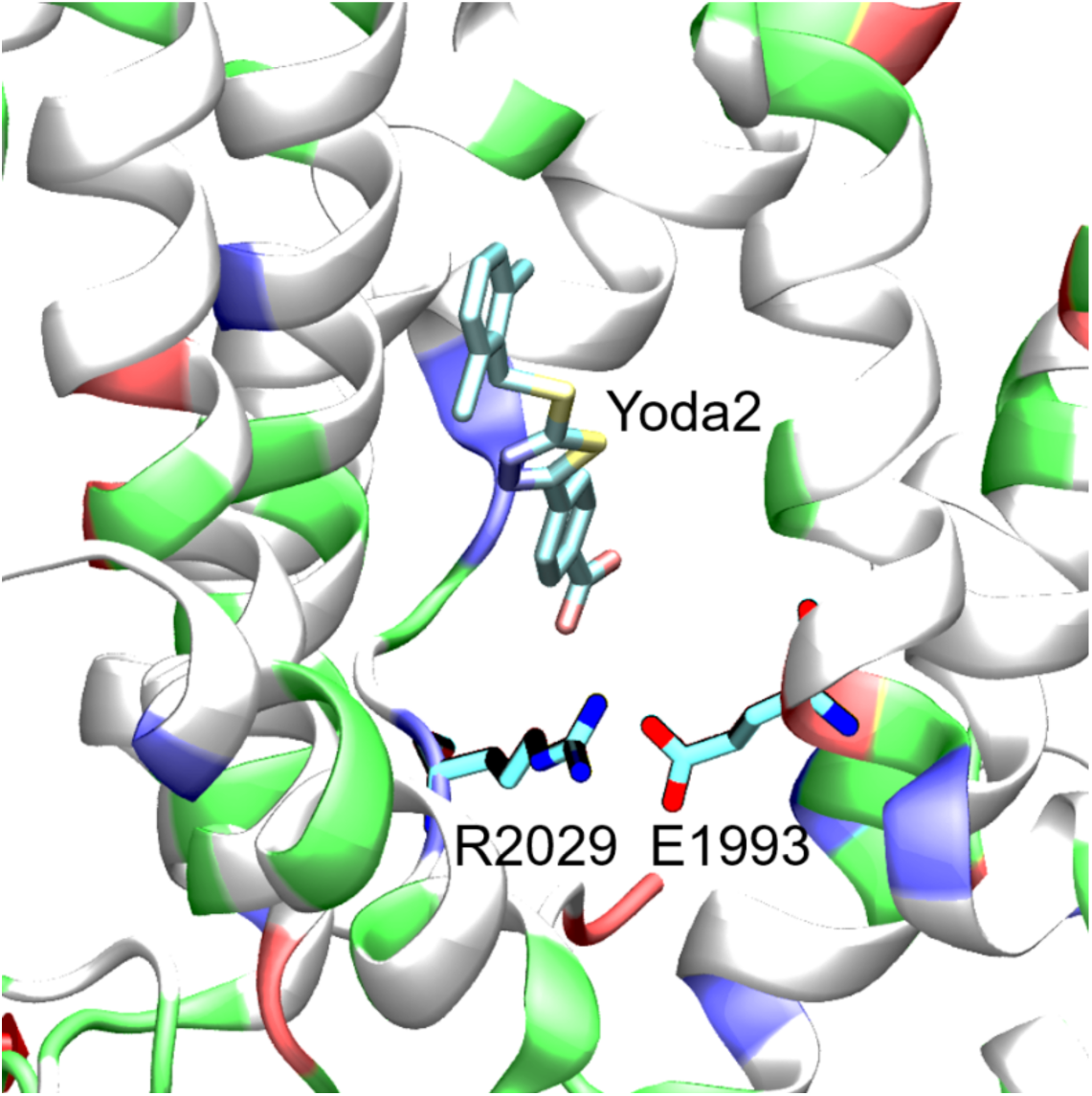
Snapshot taken from our PIEZO2 simulation Replica3 showing an electrostatic competition between Yoda2 and E1993 to form a salt-bridge with R2029.

**Figure 6 - figure supplement 4:**
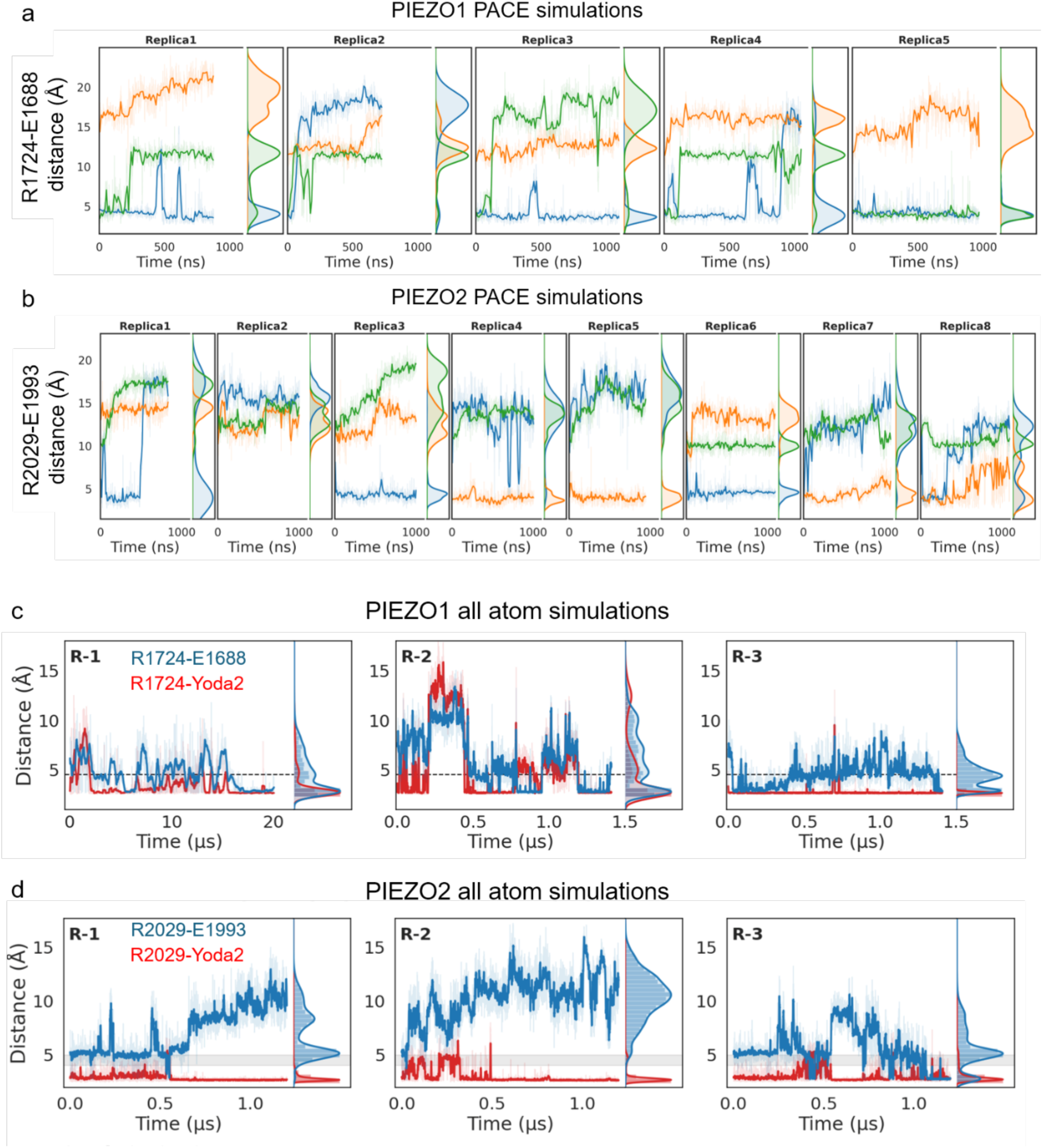
Yoda2 modulation of an endogenous salt-bridge between conserved glutamate-arginine pair in the Yoda binding site. **a-b**) Salt-bridge distances measured in all channel subunits between R1724-Cζ and E1688-Cδ for PIEZO1 (top) and between R2029-Cζ and E1993-Cδ for PIEZO2 (bottom) from previous PACE simulations (15). **c-d**) The same distances are plotted in blue for our three all-atom PIEZO1 and PIEZO2 simulation replicas in the presence of Yoda2. Salt-bridge distances between Yoda2 and R1724 (PIEZO1) and R2029 (PIEZO2) are plotted in red. Solid lines show the running average, and the panel to the right of each trace shows the distance distribution over the full trajectory.

## Notes

### Competing Interest Statement

The authors have declared no competing interest.

### Summary of Updates

We have updated all sections of the manuscripts as part of our response the the eLife reviewers. The revised texts are substantial and thus can not be listed indiividually here.

## References

1. B. Coste et al., Piezo proteins are pore-forming subunits of mechanically activated channels. Nature 483, 176–181 (2012).

2. B. Coste et al., Piezo1 and Piezo2 are essential components of distinct mechanically activated cation channels. Science 330, 55–60 (2010).

3. L. Wang et al., Structure and mechanogating of the mammalian tactile channel PIEZO2. Nature 573, 225–229 (2019).

4. K. Saotome et al., Structure of the mechanically activated ion channel Piezo1. Nature 554, 481–486 (2018).

5. Q. Zhao et al., Structure and mechanogating mechanism of the Piezo1 channel. Nature 554, 487–492 (2018).

6. Y. R. Guo, R. MacKinnon, Structure-based membrane dome mechanism for Piezo mechanosensitivity. Elife 6 (2017).

7. C. A. Haselwandter, R. MacKinnon, Piezo’s membrane footprint and its contribution to mechanosensitivity. Elife 7 (2018).

8. C. A. Haselwandter, Y. R. Guo, Z. Fu, R. MacKinnon, Elastic properties and shape of the Piezo dome underlying its mechanosensory function. Proc. Natl. Acad. Sci. U. S. A. 119, e2208034119 (2022).

9. C. A. Haselwandter, Y. R. Guo, Z. Fu, R. MacKinnon, Quantitative prediction and measurement of Piezo’s membrane footprint. Proc. Natl. Acad. Sci. U. S. A. 119, e2208027119 (2022).

10. X. Yang et al., Structure deformation and curvature sensing of PIEZO1 in lipid membranes. Nature 10.1038/s41586-022-04574-8 (2022).

11. Y. C. Lin et al., Force-induced conformational changes in PIEZO1. Nature 573, 230–234 (2019).

12. W. Jiang et al., Crowding-induced opening of the mechanosensitive Piezo1 channel in silico. Communications Biology 4, 84 (2021).

13. S. Liu et al., An intermediate open structure reveals the gating transition of the mechanically activated PIEZO1 channel. Neuron 10.1016/j.neuron.2024.11.020 (2024).

14. A. D. Ozkan, T. D. Wijerathne, T. Gettas, J. J. Lacroix, Force-induced motions of the PIEZO1 blade probed with fluorimetry. Cell Rep 42, 112837 (2023).

15. S. Li et al., A two-step clockwork mechanism opens a proteo-lipidic pore in PIEZO2. Nat. Chem. Biol. 10.1038/s41589-026-02147-8 (2026).

16. L. David et al., Piezo mechanosensory channels regulate centrosome integrity and mitotic entry. Proc. Natl. Acad. Sci. U. S. A. 120, e2213846120 (2023).

17. F. Wei et al., Deficiency of Endothelial Piezo2 Impairs Pulmonary Vascular Angiogenesis and Predisposes Pulmonary Hypertension. Hypertension 82, 583–597 (2025).

18. S. S. Ranade et al., Piezo2 is the major transducer of mechanical forces for touch sensation in mice. Nature 516, 121–125 (2014).

19. S. H. Woo et al., Piezo2 is required for Merkel-cell mechanotransduction. Nature 509, 622–626 (2014).

20. V. L. Ehlers, A. Sriram, B. A. R. Stuart, C. M. Mecca, C. L. Stucky, Sensory neuron PIEZO1 deletion inhibits dynamic light touch sensitivity in uninjured mice, prevents neuropathic light touch hypersensitivity, and drives compensatory changes in dorsal root ganglia. Pain 10.1097/j.pain.0000000000003781 (2025).

21. M. R. Servin-Vences et al., PIEZO2 in somatosensory neurons controls gastrointestinal transit. Cell 186, 3386–3399 e3315 (2023).

22. D. Florez-Paz, K. K. Bali, R. Kuner, A. Gomis, A critical role for Piezo2 channels in the mechanotransduction of mouse proprioceptive neurons. Sci. Rep. 6, 25923 (2016).

23. S. H. Woo et al., Piezo2 is the principal mechanotransduction channel for proprioception. Nat. Neurosci. 18, 1756–1762 (2015).

24. W. Sun et al., The mechanosensitive Piezo1 channel is required for bone formation. Elife 8 (2019).

25. X. Li et al., Stimulation of Piezo1 by mechanical signals promotes bone anabolism. Elife 8 (2019).

26. F. Jiang et al., The mechanosensitive Piezo1 channel mediates heart mechano-chemo transduction. Nat Commun 12, 869 (2021).

27. D. Choi et al., Piezo1 incorporates mechanical force signals into the genetic program that governs lymphatic valve development and maintenance. JCI Insight 4, 125068 (2019).

28. K. Nonomura et al., Mechanically activated ion channel PIEZO1 is required for lymphatic valve formation. Proc. Natl. Acad. Sci. U. S. A. 115, 12817–12822 (2018).

29. J. Li et al., Piezo1 integration of vascular architecture with physiological force. Nature 515, 279–282 (2014).

30. S. S. Ranade et al., Piezo1, a mechanically activated ion channel, is required for vascular development in mice. Proc. Natl. Acad. Sci. U. S. A. 111, 10347–10352 (2014).

31. S. M. Cahalan et al., Piezo1 links mechanical forces to red blood cell volume. Elife 4 (2015).

32. S. Demolombe, F. Duprat, E. Honore, A. Patel, Slower Piezo1 inactivation in dehydrated hereditary stomatocytosis (xerocytosis). Biophys. J. 105, 833–834 (2013).

33. C. Bae, R. Gnanasambandam, C. Nicolai, F. Sachs, P. A. Gottlieb, Xerocytosis is caused by mutations that alter the kinetics of the mechanosensitive channel PIEZO1. Proc. Natl. Acad. Sci. U. S. A. 110, E1162–1168 (2013).

34. L. O. Romero et al., Enhanced PIEZO1 function contributes to the pathogenesis of sickle cell disease. Proc. Natl. Acad. Sci. U. S. A. 122, e2514863122 (2025).

35. M. Szczot et al., PIEZO2 mediates injury-induced tactile pain in mice and humans. Sci. Transl. Med. 10 (2018).

36. S. E. Murthy et al., The mechanosensitive ion channel Piezo2 mediates sensitivity to mechanical pain in mice. Sci. Transl. Med. 10 (2018).

37. N. Eijkelkamp et al., A role for Piezo2 in EPAC1-dependent mechanical allodynia. Nat Commun 4, 1682 (2013).

38. V. Lukacs et al., Impaired PIEZO1 function in patients with a novel autosomal recessive congenital lymphatic dysplasia. Nat Commun 6, 8329 (2015).

39. E. Fotiou et al., Novel mutations in PIEZO1 cause an autosomal recessive generalized lymphatic dysplasia with non-immune hydrops fetalis. Nat Commun 6, 8085 (2015).

40. N. G. Amado et al., PIEZO1 loss-of-function compound heterozygous mutations in the rare congenital human disorder Prune Belly Syndrome. Nat Commun 15, 339 (2024).

41. S. Ma et al., Excessive mechanotransduction in sensory neurons causes joint contractures. Science 379, 201–206 (2023).

42. G. Haliloglu et al., Recessive PIEZO2 stop mutation causes distal arthrogryposis with distal muscle weakness, scoliosis and proprioception defects. J. Hum. Genet. 62, 497–501 (2017).

43. F. Alisch et al., Familial Gordon syndrome associated with a PIEZO2 mutation. Am J Med Genet A 10.1002/ajmg.a.37997 (2016).

44. A. A. Mahmud et al., Loss of the proprioception and touch sensation channel PIEZO2 in siblings with a progressive form of contractures. Clin. Genet. 10.1111/cge.12850 (2016).

45. M. Okubo et al., A family of distal arthrogryposis type 5 due to a novel PIEZO2 mutation. Am J Med Genet A 167A, 1100–1106 (2015).

46. M. J. McMillin et al., Mutations in PIEZO2 cause Gordon syndrome, Marden-Walker syndrome, and distal arthrogryposis type 5. Am. J. Hum. Genet. 94, 734–744 (2014).

47. B. Coste et al., Gain-of-function mutations in the mechanically activated ion channel PIEZO2 cause a subtype of Distal Arthrogryposis. Proc. Natl. Acad. Sci. U. S. A. 110, 4667–4672 (2013).

48. R. Syeda et al., Chemical activation of the mechanotransduction channel Piezo1. Elife 4 (2015).

49. G. Parsonage et al., Improved PIEZO1 agonism through 4-benzoic acid modification of Yoda1. Br. J. Pharmacol. 10.1111/bph.15996 (2022).

50. H. Jäntti et al., Microglial amyloid beta clearance is driven by PIEZO1 channels. J. Neuroinffammation 19 (2022).

51. S. Shindo et al., Piezo1 protects against inflammatory bone loss via a unique Ca-independent mechanism in osteoclasts. Front. Immunol. 16 (2025).

52. X. He et al., Yoda1 pretreated BMSC derived exosomes accelerate osteogenesis by activating phospho-ErK signaling via Yoda1-mediated signal transmission. J Nanobiotechnol 22 (2024).

53. Q. A. Meslier, R. Oehrlein, S. J. Shefelbine, Combined Effects of Mechanical Loading and Piezo1 Chemical Activation on 22-Months-Old Female Mouse Bone Adaptation. Aging Cell 24, e70087 (2025).

54. G. Silvani et al., Capillary constrictions prime cancer cell tumorigenicity through PIEZO1. Nature Communications 16 (2025).

55. W. X. Liao et al., The activation of Piezo1 channel promotes invasion and migration via the release of extracellular ATP in cervical cancer. Pathology Research and Practice 260 (2024).

56. J. U. Jung, S. Stippec, M. H. Cobb, Activation of WNK1 signaling through Piezo1. Proc. Natl. Acad. Sci. U. S. A. 122 (2025).

57. M. Lone et al., Activation of the Piezo1 channel stimulates protein kinase D and migration in human aortic endothelial cells. Am. J. Physiol. Cell Physiol. 329, C1652–C1665 (2025).

58. N. Endesh et al., Independent endothelial functions of PIEZO1 and TRPV4 in hepatic portal vein and predominance of PIEZO1 in mechanical and osmotic stress. Liver Int 43, 2026–2038 (2023).

59. M. Qiao et al., Piezo1 Channel Activators Yoda1 and Yoda2 in the Context of Red Blood Cells. Biomolecules 15 (2025).

60. T. D. Wijerathne, A. D. Ozkan, J. J. Lacroix, Yoda1’s energetic footprint on Piezo1 channels and its modulation by voltage and temperature. Proc. Natl. Acad. Sci. U. S. A. 10.1073/pnas.2202269119, e2202269119 (2022).

61. J. J. Lacroix, W. M. Botello-Smith, Y. Luo, Probing the gating mechanism of the mechanosensitive channel Piezo1 with the small molecule Yoda1. Nature Communications 9, 2029 (2018).

62. S. E. Murthy, Deciphering mechanically activated ion channels at the single-channel level in dorsal root ganglion neurons. J. Gen. Physiol. 155 (2023).

63. Z. Zhou et al., MyoD-family inhibitor proteins act as auxiliary subunits of Piezo channels. Science 381, 799–804 (2023).

64. C. Verkest, L. Roettger, N. Zeitzschel, S. G. Lechner, 3D-MINFLUX nanoscopy reveals distinct allosteric mechanisms for activation and modulation of PIEZO1 by Yoda1. Nat Commun 16, 11192 (2025).

65. T. W. Chen et al., Ultrasensitive fluorescent proteins for imaging neuronal activity. Nature 499, 295–300 (2013).

66. E. M. Mulhall, O. Yarishkin, R. Z. Hill, A. K. Koster, A. Patapoutian, The molecular basis of force selectivity by PIEZO2. Nature 10.1038/s41586-026-10182-7 (2026).

67. Y. Wang et al., A lever-like transduction pathway for long-distance chemical- and mechano-gating of the mechanosensitive Piezo1 channel. Nat Commun 9, 1300 (2018).

68. W. Jiang et al., Structural and thermodynamic framework for PIEZO1 modulation by small molecules. Proc. Natl. Acad. Sci. U. S. A. 120, e2310933120 (2023).

69. W. M. Botello-Smith et al., A mechanism for the activation of the mechanosensitive Piezo1 channel by the small molecule Yoda1. Nature Communications 10, 4503 (2019).

70. T. D. Wijerathne, A. D. Ozkan, J. J. Lacroix, Thermodynamic and structural bases for Yoda1-mediated Piezo1 activation. Biophys. J. 121, 176a (2022).

71. E. L. Evans et al., Yoda1 analogue (Dooku1) which antagonizes Yoda1-evoked activation of Piezo1 and aortic relaxation. Br. J. Pharmacol. 175, 1744–1759 (2018).

72. S. Goon et al., Exploring the Structural Attributes of Yoda1 for the Development of New-Generation Piezo1 Agonist Yaddle1 as a Vaccine Adjuvant Targeting Optimal T Cell Activation. J. Med. Chem. 67, 8225–8246 (2024).

73. H. Tang et al., Structural Modification and Pharmacological Evaluation of (Thiadiazol-2-yl)pyrazines as Novel Piezo1 Agonists for the Intervention of Disuse Osteoporosis. J. Med. Chem. 67, 19837–19851 (2024).

74. K. Xu, M. Liu, L. Wang, G. Zhan, G. He, Piezo Family: From Structure to Therapeutic Agents in Cancer. J. Med. Chem. 10.1021/acs.jmedchem.5c03200 (2026).

75. A. H. Lewis, J. Grandl, Mechanical sensitivity of Piezo1 ion channels can be tuned by cellular membrane tension. Elife 4 (2015).

76. J. Wang et al., Tethering Piezo channels to the actin cytoskeleton for mechanogating via the cadherin-beta-catenin mechanotransduction complex. Cell Rep 38, 110342 (2022).

77. A. K. Koster et al., Chemical mapping of the surface interactome of PIEZO1 identifies CADM1 as a modulator of channel inactivation. Proc. Natl. Acad. Sci. U. S. A. 121, e2415934121 (2024).

78. M. G. Dalghi, et al., Functional roles for PIEZO1 and PIEZO2 in urothelial mechanotransduction and lower urinary tract interoception. JCI Insight 6 (2021).

79. A. Savadipour et al., Membrane stretch as the mechanism of activation of PIEZO1 ion channels in chondrocytes. Proc. Natl. Acad. Sci. U. S. A. 120, e2221958120 (2023).

80. S. Akita et al., Increased expression of the PIEZO2 mechanoreceptor in fibroblasts and endothelial cells within the lymphatic and vascular vessels of keloids. J. Pathol. 267, 105–119 (2025).

81. H. M. Nguyen, K. Y. Huang, Y. J. Cui, H. Shim, H. Wulff, Off-target effects of Piezo1 agonists on macrophages. Biophys. J. 124 (2025).

82. S. Pourteymour et al., PIEZO1 targeting in macrophages boosts phagocytic activity and foam cell apoptosis in atherosclerosis. Cell Mol Life Sci 81, 331 (2024).

83. Y. Xiao et al., ZnCe-LDO Nanozyme-Based Multifunctional Hydrogel Promotes Bone Regeneration by Inflammatory Macrophage Reprogramming and Piezo1 Activation. ACS Nano 19, 32606–32628 (2025).

84. S. Shindo et al., Piezo1 protects against inflammatory bone loss via a unique Ca(2+)-independent mechanism in osteoclasts. Front. Immunol. 16, 1661538 (2025).

85. N. G. dela Paz, J. A. Frangos, Yoda1-induced phosphorylation of Akt and ERK1/2 does not require Piezol activation. Biochemical and Biophysical Research Communications 497, 220–225 (2018).

86. C. Alcaino, K. Knutson, P. A. Gottlieb, G. Farrugia, A. Beyder, Mechanosensitive ion channel Piezo2 is inhibited by D-GsMTx4. Channels (Austin) 11, 245–253 (2017).

87. C. Bae, F. Sachs, P. A. Gottlieb, The mechanosensitive ion channel Piezo1 is inhibited by the peptide GsMTx4. Biochemistry 50, 6295–6300 (2011).

88. K. L. Ostrow et al., cDNA sequence and in vitro folding of GsMTx4, a specific peptide inhibitor of mechanosensitive channels. Toxicon 42, 263–274 (2003).

89. T. D. Wijerathne, A. Bhatt, W. Jiang, Y. L. Luo, J. J. Lacroix, Mammalian PIEZO channels rectify anionic currents. Biophys. J. 10.1016/j.bpj.2024.11.010 (2024).

